# Flux through lipid synthesis dictates bacterial cell size

**DOI:** 10.1101/092684

**Authors:** Stephen Vadia, Jessica L. Tse, Jue D. Wang, Petra Anne Levin

## Abstract

Nutrients—and by extension biosynthetic capacity—positively impact cell size in organisms throughout the tree of life. In bacteria, cell size is reduced three-fold in response to nutrient starvation or accumulation of the alarmone ppGpp, a global inhibitor of biosynthesis. However, whether biosynthetic capacity as a whole determines cell size or if particular anabolic pathways are more important than others remains an open question. Utilizing a top-down approach, here we identify flux through lipid synthesis as the primary biosynthetic determinant of *Escherichia coli* cell size. Altering flux through lipid synthesis recapitulated the impact of altering nutrients on cell size and morphology, while defects in other biosynthetic pathways either did not impact size or altered size in a lipid-dependent manner. Together our findings support a model in which lipid availability dictates cell envelope capacity and ppGpp functions as a linchpin linking surface area expansion with cytoplasmic volume to maintain cellular integrity.

## Introduction

Cell size control is a fundamental aspect of biology. In multicellular organisms differences in cell size frequently dictate developmental fate, and defects in cell size are associated with cancer and other disease states (Ginzberg et al., 2015). In single celled organisms defects in cell size can be catastrophic, leading to failures in chromosome partitioning, bisection of genomic material during cytokinesis, and cell death (Hill et al., 2013; Rowlett and Margolin, 2015; Weart et al., 2007).

With the exception of asymmetric division and the formation of syncytia, cell size is dictated by two parameters: growth rate and cell cycle progression. Assuming growth rate remains constant, increasing the rate of cell cycle progression results in reductions in cell size. Conversely, maintaining the rate of cell cycle progression but increasing growth rate results in increases in cell size. Balancing these two parameters ensures cell size is maintained, an idea supported by the recent discovery that bacterial cells cultured under steady state conditions add a constant volume (Δ) prior to division regardless of their size at birth (Amir, 2014; Campos et al., 2014; Iyer-Biswas et al., 2014; Taheri-Araghi et al., 2015). Follow-up work indicates that deviations in birth size between *E. coli* cells in a clonal population are a consequence of stochastic differences in growth rate (Wallden et al., 2016).

Research on cell size homeostasis has historically focused on the role of the cell cycle. Most famously, the *wee1* kinase from *Schizosaccharomyces pombe* influences the timing of mitosis to maintain cell size, a function conserved in its many eukaryotic homologs (Kellogg, 2003; Wood and Nurse, 2015). In bacteria including the evolutionarily distant model organisms *Escherichia coli* and *Bacillus subtilis*, cell size is coupled with carbon availability in part via the UDP-glucose-dependent activation of a division inhibitor that delays the onset of cytokinesis to increase cell length (Hill et al., 2013; Weart et al., 2007). Growth-dependent accumulation of the DNA replication initiation factor DnaA has also been implicated in cell size homeostasis under steady-state conditions through its impact on the timing and frequency of replication initiation (Lobner-Olesen et al., 1989; Wallden et al., 2016).

Although it has received considerably less attention, several lines of evidence support the idea that biosynthetic capacity, a product of nutrient availability, also serves as a primary determinant of cell size. Most important among these is the positive correlation between nutrient availability and cell size that exists throughout the tree of life. Drosophila cell and body size scale with nutrients, as does the size of baker’s yeast, *Saccharomyces cerevisiae*, and fission yeast, *Schizosaccharomyces pombe* (Davie and Petersen, 2012; Edgar, 2006). The relationship between cell size, nutrient availability, and growth rate is linear in *Salmonella, E. coli* and *B. subtilis.* In all three bacterial species, organisms cultured under nutrient-rich conditions are up to three times the volume of those cultured under nutrient-poor conditions, a size increase too great to be accounted for solely by UDP-glucose-dependent division inhibition (Hill et al., 2013; Pierucci, 1978; Sargent, 1975; Schaechter et al., 1958; Weart et al., 2007). Significantly, Δ, the volume of material that *E. coli* cells add each generation, increases in carbon rich conditions and decreases in carbon poor conditions, suggesting that changes in biosynthetic capacity shift the balance between growth rate and the rate of cell cycle progression (Campos et al., 2014; Taheri-Araghi et al., 2015).

In further support of biosynthetic capacity as an important determinant of cell size, accumulation of guanosine tetraphosphate (ppGpp), an inhibitor of a host of biosynthetic pathways, is negatively correlated with bacterial size. Increasing intracellular ppGpp concentration directly via overexpression of the ppGpp synthase RelA reduces the growth rate and size of *E. coli* cultured in nutrient rich medium (Schreiber et al., 1995; Tedin and Bremer, 1992). Indirectly stimulating ppGpp accumulation via amino acid starvation or by limiting fatty acid synthesis, similarly leads to reductions in both growth rate and cell size (Tehranchi et al., 2010; Yao et al., 2012a).

Because ppGpp is a global regulator of biosynthesis in bacteria (Liu et al., 2015), however, it is impossible to know if ppGpp-dependent reductions in size are the result of changes in biosynthetic capacity as a whole, or if particular anabolic pathways play a more important role than others in modulating cell size. Thus, while amino acid starvation or limitations in fatty acid synthesis may directly impact cell size, it is equally likely that they are affecting size indirectly via increases in ppGpp concentration and the concomitant global down-regulation of biosynthetic capacity. This precise conundrum was noted by Yao *et al.* in their 2012 paper reporting a reduction in *E. coli* size associated with defects in a gene required for an early step in fatty acid synthesis (Yao et al., 2012a).

To illuminate the role of biosynthetic capacity in cell size control, we employed a top-down strategy to assess the contribution of three major anabolic pathways to *E. coli* morphology and growth rate. Our findings unequivocally and for the first time establish flux through lipid synthesis as a primary, ppGpp-independent determinant of bacterial cell size. Not only did increasing flux through lipid synthesis lead to dramatic ppGpp-independent increases in cell size, defects in other major pathways impacted by ppGpp had either no significant impact on size (RNA synthesis) or reduced size in a lipid-dependent manner (protein synthesis). Importantly, increases in flux through lipid synthesis significantly increased both cell length and width, almost perfectly recapitulating the effect of nutrients on cell morphology in both magnitude and kind. The positive relationship between lipid synthesis and cell width distinguishes it from cell cycle progression, changes in which almost exclusively impact cell length (Harris and Theriot, 2016; Pritchard et al., 1978; Young, 2010).

Taken together our findings support a novel “outside-in” model in which size is dictated by the capacity of the cell envelope, which is itself a product of nutrient-dependent changes in lipid synthesis and availability. This lipid-centric view has implications for essentially all aspects of bacterial physiology that are sensitive to changes in nutrient availability. In particular, changes in cell envelope biogenesis must be coordinated with synthesis of cytoplasmic material to ensure that cytoplasmic volume does not exceed plasma membrane capacity. Our data suggest that ppGpp plays a central role in this process, tethering lipid synthesis to cytoplasmic aspects of anabolic metabolism to preserve cell envelope integrity in the face of a rapidly changing nutritional landscape.

## Results

### RNA, protein and fatty acid synthesis differentially impact cell size

To identify the major biosynthetic pathways underlying cell size control, we utilized a top-down approach, modifying the growth rate of *E. coli* cultured in nutrient-rich medium (LB + 0.2% glucose; LB-glc) with subinhibitory concentrations of antibiotics targeting RNA (rifampicin), protein (chloramphenicol), and fatty acid (cerulenin) synthesis. Synthesis of RNA, protein, and fatty acids is nutrient-dependent, and all three pathways are down-regulated in response to increases in ppGpp levels, either directly (RNA), or indirectly (protein and fatty acid synthesis) (Heath et al., 1994; Podkovyrov and Larson, 1996; Schreiber et al., 1991; Svitil et al., 1993). In light of the well-documented association between defects in DNA replication and activation of both SOS-dependent and -independent modes of division inhibition, we elected not to investigate the impact of reductions in DNA synthesis on size (Arjes et al., 2014; Cambridge et al., 2014; D’Ari and Huisman, 1983; Huisman and D’Ari, 1981; Huisman et al., 1984). We similarly chose not to examine the consequence of defects in cell wall biogenesis on cell size, as treatment with antibiotics targeting peptidoglycan synthesis disproportionally effect cell division, leading to extensive filamentation even at low concentrations (Burke et al., 2013; Fredborg et al., 2015; Yao et al., 2012b).

To assess the impact of limitations in RNA, protein, or fatty acid synthesis on cell size, BH330 cells [MG1655 *P_lac_∷gfp-ftsZ* (*bla*) (Hale and de Boer, 1999)] were first cultured to early log phase in LB-glc, then back-diluted to an OD600 of 0.005 into fresh medium with subinhibitory concentrations of the appropriate antibiotic and 1 mM IPTG to induce expression from *P_lac_-gfp-ftsZ*, and cultured for ~ 6 generations (OD600 ~ 0.3) prior to being fixed, imaged, and measured. Notably, we did not detect any significant differences in growth rate or cell size if chloramphenicol or cerulenin treated cells were back-diluted a second time into fresh medium and cultured for an additional 6 generations at the same drug concentration. However, in the presence of the highest concentration of rifampicin (6 μg/ml), cells were unable to reach a substantial OD600 upon back-dilution, suggesting that robust transcription is a requirement for exit from early log growth (Table S3). As a benchmark, we also modified growth by culturing cells in six different media: LB-glc, LB, and AB minimal medium ± 0.5% casamino acids with 0.2% glucose or 0.4% succinate (AB-caa-glc, AB-caa-succ, AB-glc, AB-succ). Measurements from ≥400 cells from ≥3 biological replicates were used to generate single data points and histograms for each condition. Cell length and width analysis was performed using the Image J plug-in, Coli-Inspector (Vischer et al., 2015).

Consistent with previous studies, reductions in nutrient availability led to decreases in both growth rate and size (Figures 1 and S1) (Pierucci, 1978; Sargent, 1975; Schaechter et al., 1958). *E. coli* cultured in LB-glc grew to a mean length (*l*) of 3.76 μm, mean width (*w*) of 1.07 μm and mean square area (*a*) of 4.11 μm^2^, while those cultured in minimal succinate, the poorest carbon source, were 48% smaller by area (*a* = 2.07 μm^2^) in addition to being shorter and thinner (*l* =2.63 μm, *w* = 0.79 μm).

**Figure 1.**
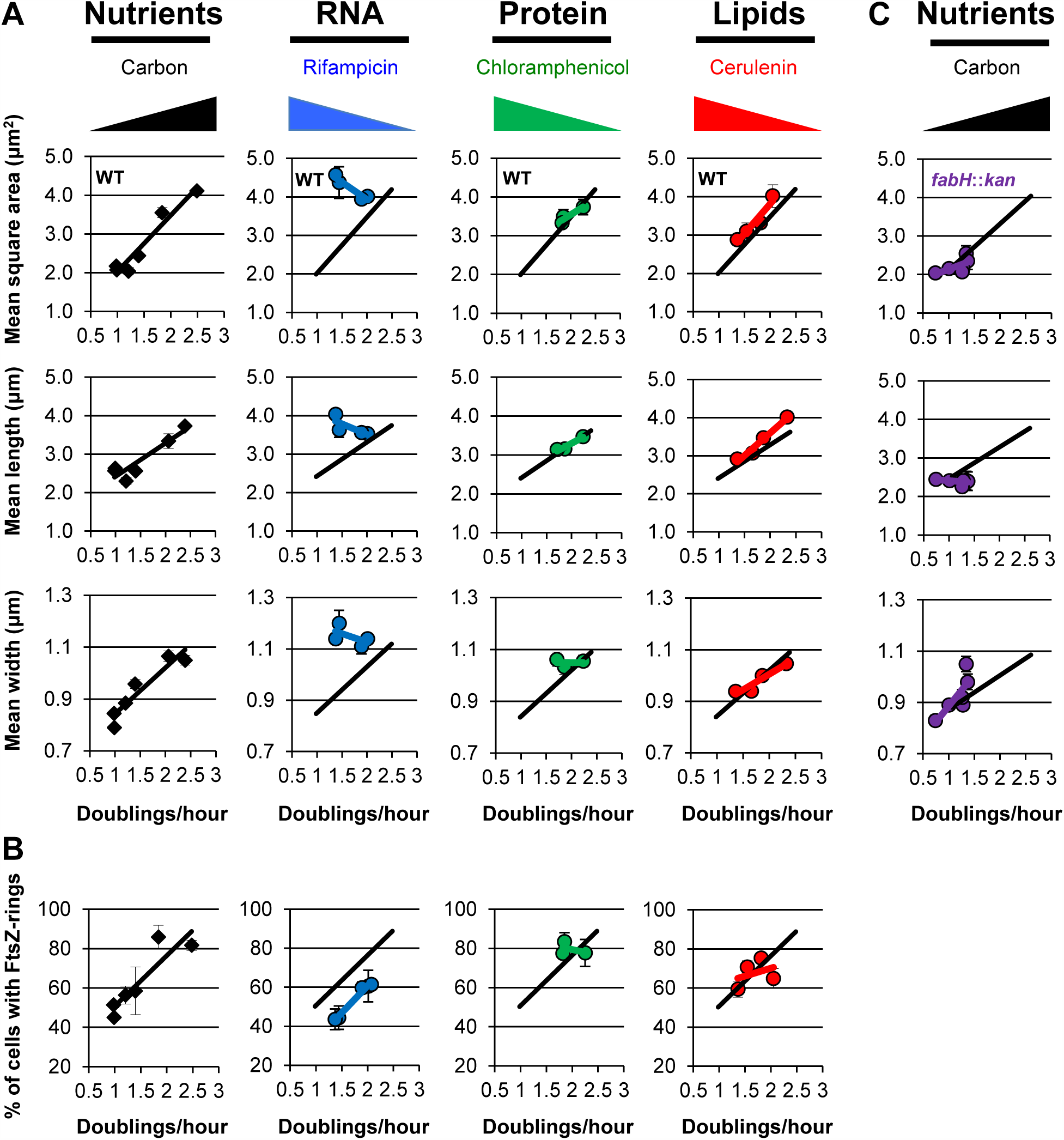
RNA, protein and fatty acid synthesis differentially impact cell size. (A) Plots of mass doublings/hour versus mean square area, mean length and mean width for cells cultured in different carbon sources, or cultured in LB-glc and treated with subinhibitory concentrations of antibiotics inhibiting RNA (rifampicin), protein (chloramphenicol), or fatty acid synthesis (cerulenin). (B) Plots of doubling time versus percent of cells with FtsZ rings for the above conditions. (C) Mass doublings/hour versus mean square area for cells with loss-of-function mutations in *fabH*, whose product catalyzes the conversion of acetyl-CoA and malonyl-acyl-carrier-protein to acetoacetyl-acyl-carrier-protein in one of the first steps in fatty acid synthesis. All data points indicate averages from n ≥3 experiments, ≥100 cells measured per experiment. Error bars = standard error of the mean (SEM). Black lines indicate the line of best fit of data from cells cultured in different carbon sources. Blue, green, red, and purple lines indicate lines of best fit for respective treatments. Wedges indicate increasing carbon availability or antibiotic concentration for respective conditions.

Of the three antibiotics, only cerulenin, an inhibitor of the β-ketoacyl-ACP synthase I, FabB, reduced both growth rate and size in a manner that closely mimicked effects of nutrient limitation (Figures 1 and S1A; For reference the carbon curve is depicted as a black line in the plots of cell size versus growth rate following antibiotic treatment). In the presence of 70 μg/ml of cerulenin, cells cultured in LB-glc exhibited a growth rate and size similar to that of untreated cells cultured in AB-caa-glc (LB-glc + cer: mean mass doubling time/hour (τ) = 1.36 d/h, *l* = 2.93 μm, *w* = 0.95 μm, *a* = 2.88 μm^2^; AB-caa-glc: τ = 1.4 d/h, *l* = 2.56 μm, *w* = 0.96 μm, *a* = 2.44 μm^2^). This reduction in size is similar to that previously reported for cerulenin treated cells (Yao et al., 2012a). While treatment with between 0.5 μg/ml and 1.5 μg/ml of chloramphenicol reduced cell length to 3.14 μm, width remained essentially constant (~1.0 to 1.1 μm). Inhibiting transcription with rifampicin dramatically reduced growth rate but did not reduce cell size at even the highest concentration of antibiotic (6 μg/ml). The 10% reduction we observed in chloramphenicol-treated cell size is similar to that previously reported by the Hwa and Theriot labs (Basan et al., 2015a; Basan et al., 2015b; Harris and Theriot, 2016). The size and growth rate of BH330 cells cultured under different nutrient conditions were essentially identical to the size and growth rate of the wild type MG1655 parental strain cultured under the same conditions, indicating that expression of *P_lac_∷gfp-ftsZ* did not significantly impact cell growth or physiology (Figure S1B).

While the three antibiotics differentially impacted cell size, none of them significantly disrupted the relationship between growth rate and cell cycle progression. Assembly of the tubulin-like protein FtsZ into a ring-like structure at the nascent division site is an essential phase of cytokinesis in bacteria. The FtsZ ring serves as a framework for assembly of the division machinery and is a useful marker for cell cycle progression (Adams and Errington, 2009; den Blaauwen et al., 1999). While 80-90% of cells cultured in nutrient rich conditions have FtsZ rings, rings are visible in only 40-50% of their slow growing counterparts cultured in nutrient poor medium, consistent with a reduced rate of cell cycle progression (den Blaauwen et al., 1999; Weart and Levin, 2003).

Taking advantage of the *P_lac_∷gfp-ftsZ* construct in BH330 cells, we determined that the proportion of cells with distinct FtsZ rings (visualized as two parallel fluorescent spots at midcell, a band across the width of the cell, or as a single point in a constricting cell) remained inversely proportional to mass doubling time, regardless of the method (nutrient depletion or antibiotic treatment) used to reduce growth rate (Figure 1B). Although depressed relative to other growth conditions - potentially due to reductions in expression of both native *ftsZ* and *P_lac_∷gfp-ftsZ* - the frequency of FtsZ rings remained inversely proportional to mass doubling time in rifampicin treated cells (Figure 1B).

### Cells lacking *fabH* adjust their width in response to nutrient availability

The dose-dependent effect of cerulenin on cell size suggests a proportional relationship between size and flux through fatty acid synthesis. To test this idea we took advantage of previous work from the Ruiz laboratory, indicating that *E. coli* cells defective in *fabH*, encoding the enzyme responsible for catalyzing the initiating condensation reaction in fatty acid synthesis (β-ketoacyl-acyl carrier protein synthase III), are viable due to the presence of an as yet uncharacterized bypass mechanism that permits growth in the absence of what was previously assumed to be an essential gene (Yao et al., 2012a).

Reasoning that if size is in fact proportional to flux through fatty acid synthesis, as our cerulenin data suggest, the linear relationship between nutrient availability and cell size should be partially retained in a *fabH* mutant due to the presence of the putative bypass mechanism. In support of a direct relationship between flux through fatty acid synthesis and cell size, *fabH* mutants increased in size in response to increases in carbon availability, although not to the same extent or in the same manner as wild-type *E. coli. fabH* mutants cultured in the poorest medium (AB-succ) (τ = 0.74 d/h), had a mean square area (a) of 2.04 μm^2^, those cultured in intermediate conditions, AB-caa-glc (τ = 1.28 d/h) and AB-caa-succ (τ = 1.00 d/h), both had mean square areas of 2.15 μm^2^, and those cultured in the richest medium, LB-glc (τ = 1.34 d/h), a mean square area of 2.54 μm^2^ (Figure 1C). Interestingly, length remained relatively constant across growth conditions (LB-glc *l* = 2.42 μm; AB-succ *l* = 2.45 μm) but width changed substantially (LB-glc *w* = 1.05 μm; AB-succ *w* = 0.83).

We note that our finding that *fabH* mutants maintain a positive relationship between size and nutrient availability runs counter to those of Yao *et al.*, who reported that *fabH* mutants are compromised in their ability to modulate size in response to nutrients. In their hands *fabH* mutant length and width were essentially identical in rich or minimal media (LB and M63 minimal salts + 0.2% glucose) [Table 4 (Yao et al., 2012a)]. We attribute this discrepancy to the broad range of carbon sources we employed, which in turn allowed for a broader range in flux through fatty acid synthesis.

### Protein synthesis-dependent changes in cell size are lipid-dependent

Our observation that treatment with chloramphenicol resulted in a modest but significant reduction in cell size (Figures 1 and S1), suggested that protein synthesis is either a direct determinant of cell size or, alternatively, that it indirectly impacts size via secondary effects on fatty acid synthesis. To resolve this issue, we took advantage of *E. coli’s* ability to incorporate exogenous long chain fatty acids, such as oleic acid, into its plasma membrane (Black and DiRusso, 2003; Cronan, 2014). Briefly, we cultured BH330 cells in LB-glc with 1.5 μg/ml of chloramphenicol, in the presence or absence of exogenous oleic acid. As controls we also added oleic acid to cells cultured in the presence of 70 μg/ml cerulenin and to untreated cells.

Supporting a primary role for lipids in cell size control, the addition of exogenous long chain fatty acids reversed both the growth rate and size defect of cerulenin and chloramphenicol-treated cells (Figure 2A and B; Table S4). As expected, cerulenin-treated cells increased substantially in size and growth rate in the presence of 10 μg/ml of oleic acid. Strikingly, and in agreement with the idea that chloramphenicol impacts size indirectly via reductions in cellular levels of key enzymes involved in fatty acid synthesis, exogenous oleic acid partially corrected the growth rate and cell size defects of chloramphenicol-treated cells, which were 10% larger and grew 16% faster after 5-6 generations in the presence of 10 μg/ml oleic acid than in its absence (+ oleic acid: *a* = 3.70 ± 0.16 μm^2^, τ = 2.09 d/h; - oleic acid: *a* = 3.35 ± 0.15 μm^2^ and τ = 1.82 d/h) (Figure 2B). Cell width remained essentially unchanged in the presence and absence of oleic acid (*w* = 1.10 ± 0.03). Oleic acid had no impact on bacteria in the absence of antibiotics, suggesting they were at steady state with regard to synthesis and incorporation of fatty acids (Figure S1C).

**Figure 2.**
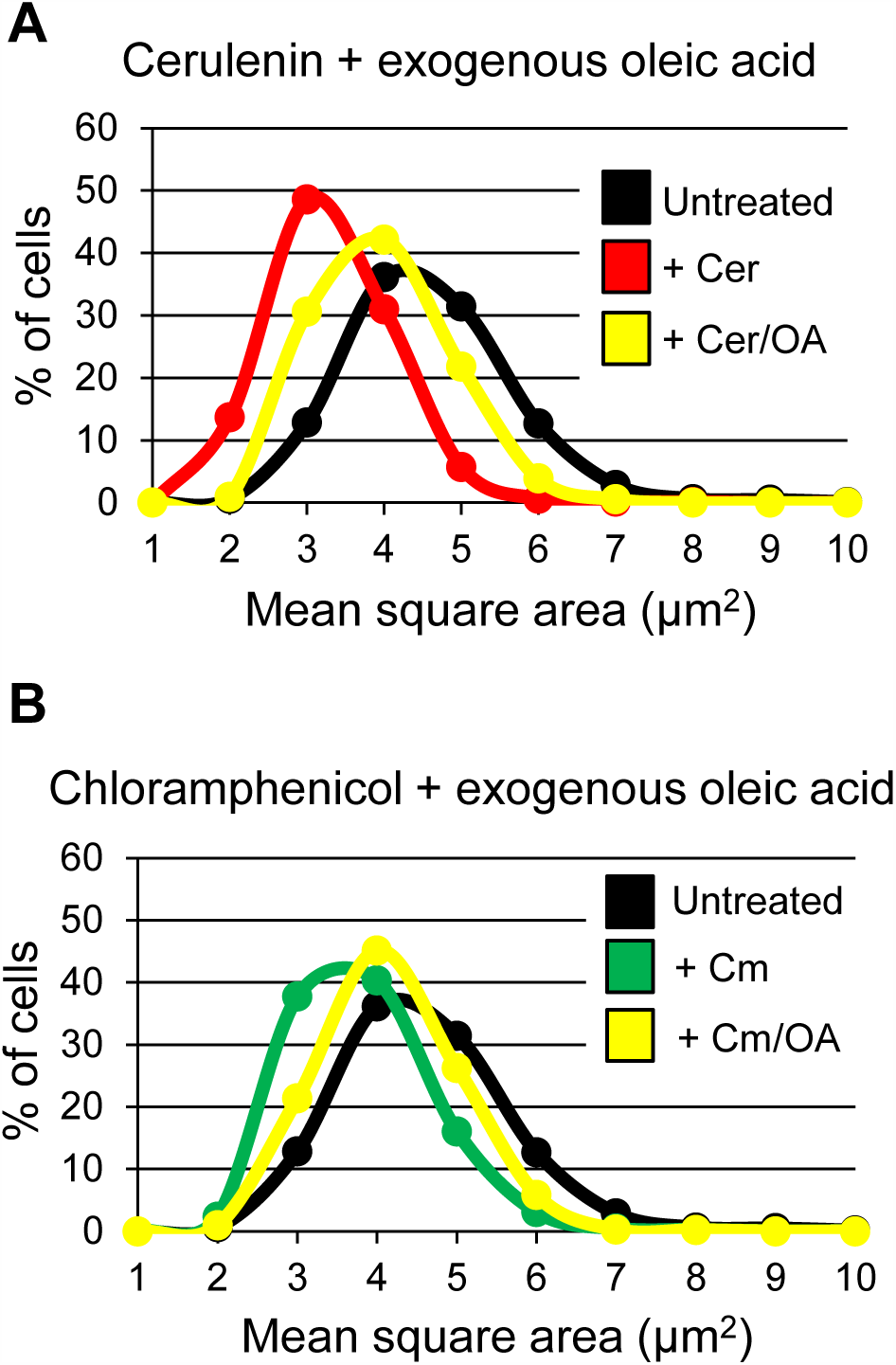
Protein-dependent changes in cell size are lipid-dependent. Cell size distributions following inhibition of fatty acid synthesis (A) or protein synthesis (B) in the presence or absence of exogenous oleic acid. Cer = 70 μg/ml cerulenin. Cm = 1.5 μg/ml chloramphenicol. OA = 10 μg/ml oleic acid. Distribution curves compiled from n ≥3 experiments for each condition, ≥100 cells measured in each experiment.

### Cell size scales with lipid synthesis independent of ppGpp

A significant limitation of the preceding experiments as well as previous work investigating the relationship between fatty acid synthesis and cell size, has been the use of mutations or antibiotics that interfere with the earliest steps in fatty acid synthesis and lead to a four-to five-fold increase in intracellular ppGpp levels due to reductions in SpoT hydrolase activity (Battesti and Bouveret, 2006; Seyfzadeh et al., 1993). ppGpp in turn down-regulates transcription of a host of genes including those encoding ribosomal RNAs, and enzymes essential for fatty acid and phospholipid synthesis (Heath et al., 1994; Liu et al., 2015). Inhibiting early steps in fatty acid biosynthesis may thus impact size indirectly, via a ppGpp-dependent but fatty acid-independent mechanism.

To clarify the relationship between lipid synthesis and cell size we therefore sought to manipulate lipid pools through alternative means: increasing synthesis of the entirety of genes required for fatty acid synthesis via induction of the transcriptional activator FadR. FadR functions as both a transcriptional activator, driving expression of genes required for fatty acid synthesis, and as a repressor, dampening expression of genes involved in the breakdown of fatty acids through beta-oxidation in response to nutritional stress (My et al., 2013). Overproduction of FadR leads to the bulk accumulation of long chain fatty acids in wild-type cells (Zhang et al., 2012). To overproduce FadR, we employed plasmid *pE8k-fadR* (hereafter *pFadR*) encoding an inducible/repressible allele of *fadR* (the kind gift of Dr. Fuzhong Zhang).

Consistent with a positive relationship between fatty acid synthesis and cell size, induction of *fadR* led to a dose-dependent increase in cell size (Figure 3A and B; Table S5). Even in the absence of inducer, the mean square area of MG1655 cells bearing *pFadR* was ~ 20% higher than MG1655 without the plasmid (*pFadR a* = 4.60 μm^2^; MG1655 *a* = 3.77 μm^2^) exhibiting increases in both length and width (*pFadR l* = 3.99 μm, *w* = 1.14 μm; MG1655 *l* = 3.53 μm, *w* = 1.07 μm). At 10μM IPTG, the highest concentration of inducer, *pFadR* cells were nearly twice the size of those from the parental strain (*pFadR* 10μM IPTG *a=* 7.15 μm^2^, *l* = 5.67, *w* = 1.23). Importantly, overproduction of FadR dramatically increased not only average cell size, but also shifted the entire cell size distribution towards larger cells (Figure 3C).

**Figure 3.**
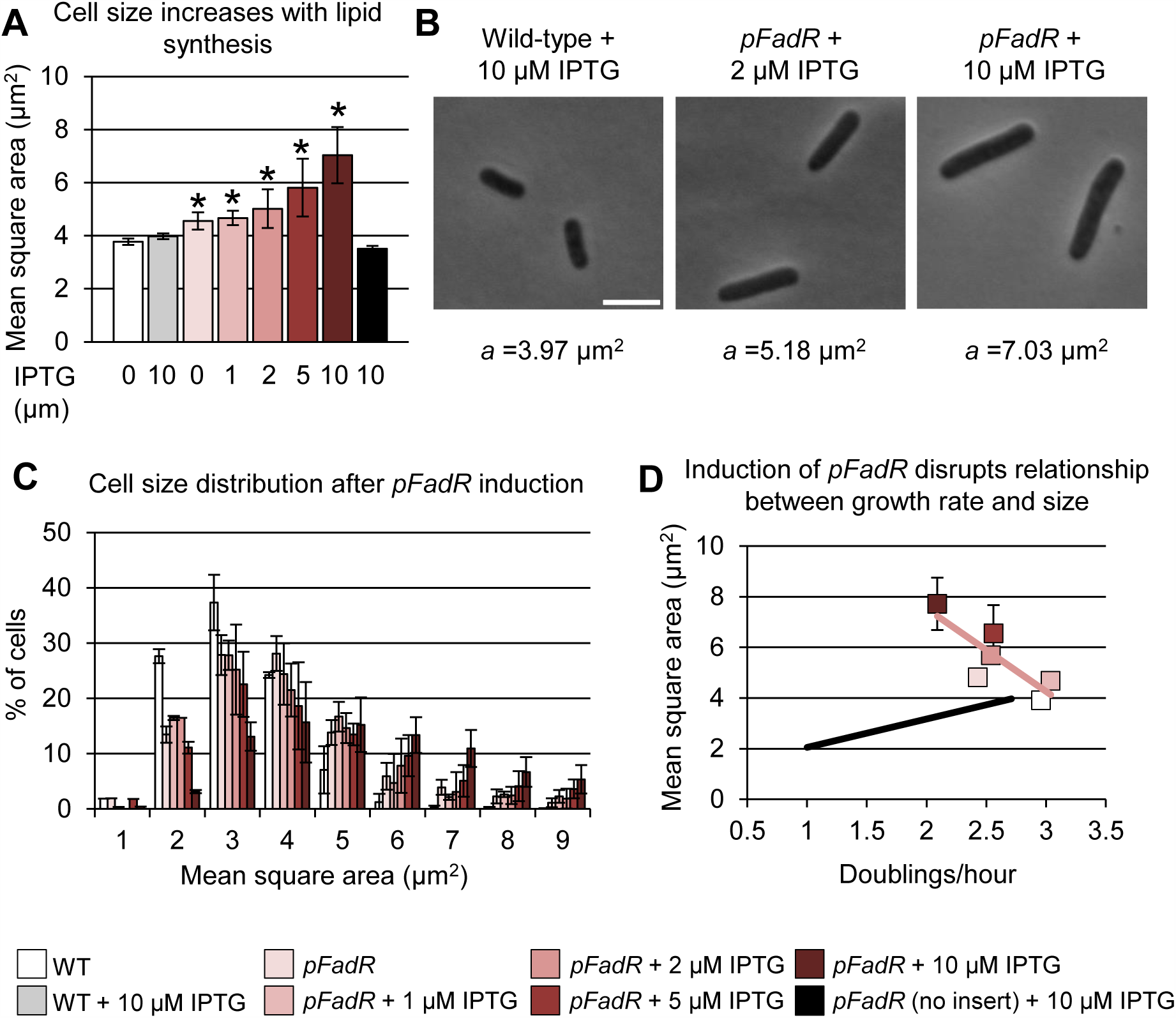
Cell size scales with lipid availability. (A) Overproduction of the transcriptional activator of fatty acid synthesis, FadR, increases cell size. Asterisks indicate a significant difference compared to wild-type MG1655. * p < 0.05 by t-test. (B) Representative phase contrast images of *E. coli* MG1655 or *MG1655-pFadR.* Scale bar = 5 μm. (C) Cell size histogram showing cell size distribution versus induction of*pFadR.* (D) Plot of cell size versus growth rate following induction of *pFadR.* Black line indicates best fit data from MG1655 cultured in different carbon sources (reproduced from Figure 1C). Red line indicates best fit data for MG1655-*pFadR*. Means are compiled from n ≥3 experiments for each condition, ≥ 100 cells measured in each experiment. Error bars = SEM. Color key indicates different IPTG concentrations and applies to panels A, C and D.

Intriguingly, induction of *fadR* inverted the relationship between size and growth rate. While *pFadR* cells were approximately twice as large as the parental strain in the presence of maximal amounts of inducer, they grew nearly 30% slower (*pFadR* 10 μM IPTG τ = 2.09 d/h; MG1655 τ = 2.88 d/h) (Table S5). This reduction in growth rate is most likely a consequence of reductions in other aspects of anabolic metabolism due to increased carbon flux through lipid synthesis. Importantly, induction of *fadR* did not significantly impact the ratio of ppGpp to GTP, consistent with a model in which fatty acid synthesis influences size independent of the alarmone (Figure S2A).

### Flux through lipid synthesis impacts cell size downstream of ppGpp

To confirm if lipid synthesis impacts size downstream of ppGpp as our *fadR* overexpression data suggested, we next assessed the ability of FadR to overcome the size defect associated with increases in intracellular ppGpp levels. ppGpp together with the RNA polymerase binding protein DksA, inhibits transcription of the genes required for fatty acid synthesis as well as *fadR* (My et al., 2013). If ppGpp mediates size through down-regulation of fatty acid synthesis, overproduction of FadR should increase the size of cells overexpressing ppGpp. Alternatively, if ppGpp impacts size downstream of fatty acid synthesis, overproduction of FadR should have little impact on the size of cells overexpressing ppGpp.

For these experiments we utilized a plasmid (*pALS10* hereafter *pRelA*; the kind gift of Dr. Charles Rock) encoding a copy of the ppGpp synthase *relA* under the control of an IPTG inducible promoter (Svitil et al., 1993). Consistent with previous studies, ppGpp had a negative impact on cell size. In the absence of inducer, *pRelA* cells were on average 25% smaller than the MG1655 parental strain (*pRelA a* = 2.94 μm^2^; *l* = 3.08 μm, *w* = 0.96 μm) (Figure 4A; Table S5). The addition of 10μM IPTG further reduced *pRelA* cells to ~55% of wild-type size (*pRelA* + 10 μM IPTG *a* = 2.20 μm^2^; *l* = 2.34 μm, *w* = 0.94 μm). *pRelA* also led to a marked increase in mass doubling time, consistent with reports indicating that even modest increases in intracellular ppGpp levels have a strong, negative impact on growth (*pRelA* τ = 2.09; *pRelA* + 10 μM IPTG τ = 1.46; MG1655 τ = 2.64) (Sarubbi et al., 1988).

**Figure 4.**
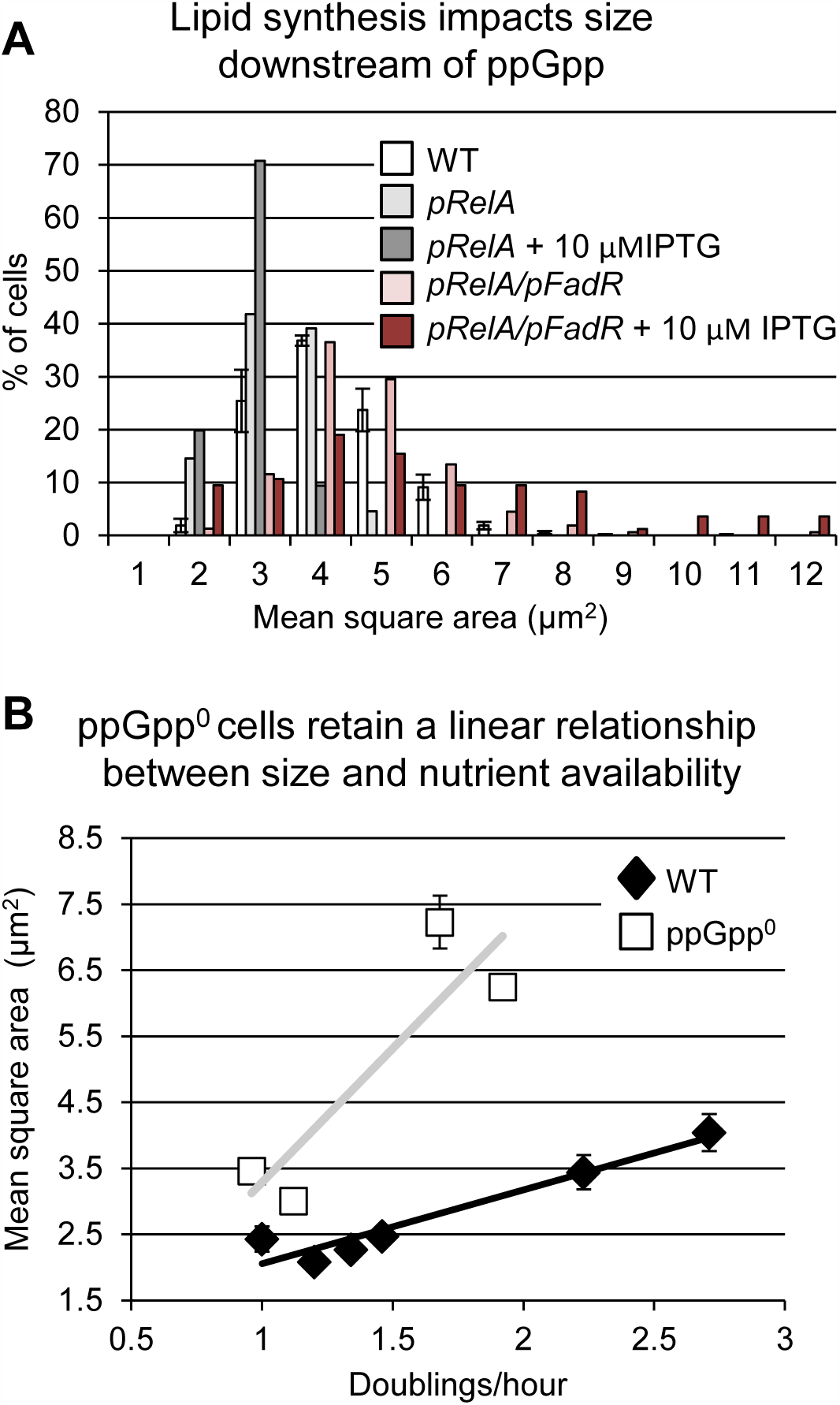
Lipid synthesis and nutrient availability impact size independent of ppGpp. (A) Histogram of cells engineered to overproduce the ppGpp synthase RelA alone and in combination with the transcriptional activator FadR in response to 10 μM IPTG (B) Plot of cell size versus growth rate of wild-type MG1655 and ppGpp^0^ strains cultured in different carbon sources: LB-glc, LB, AB-caa-glc, AB-caa-succ, for both strains as well as AB-glc and AB-succ for the wild-type strain. (The ppGpp^0^ strain cannot grow without the addition of casamino acids). Black and grey lines indicate lines of best fit for wild-type and ppGpp^0^ data sets, respectively. Compiled from n ≥3 experiments for each condition, ≥100 cells measured in each experiment. Error bars = SEM.

To ascertain if lipid synthesis impacts size downstream of ppGpp, we next generated a strain carrying both *pRelA* and *pFadR*. Because *relA* and *fadR* are both under the control of an IPTG-inducible promoter, FadR and ppGpp levels should both rise in the presence of IPTG. Consistent with a model in which ppGpp impacts size directly via down-regulation of fatty acid synthesis, cells harboring both *pRelA* and *pFadR*, were larger than cells carrying *pRelA* alone regardless of the presence of inducer (*pRelA/pFadR* - inducer *a* = 4.15 μm^2^; *l* = 3.89 μm, *w* = 1.07 μm; *pRelA/pFadR* 10 μM IPTG *a* = 4.76 μm^2^; *l* = 4.29 μm, *w* = 1.09 μm) (Figure 4A; Table S5). Co-induction of *fadR* and *relA* did not substantially alter the growth rate of *pRelA/pFadR* cells (*pRelA/pFadR* - inducer τ = 1.92 d/h; *pRelA/pFadR* + 10 μM IPTG τ = 1.71 d/h), nor did it significantly impact the ppGpp to GTP ratio (Figure S2B).

### The linear relationship between cell size and nutrient availability is retained in the absence of ppGpp

If flux through fatty acid synthesis is indeed a primary determinant of cell size as the above data suggest, *E. coli* should retain a linear relationship between size and nutrient availability regardless of the presence of ppGpp. To test this possibility, we examined the growth rate and size of *E. coli* defective in ppGpp synthesis (ppGpp^0^) cultured under four different carbon conditions (LB-glc, LB, AB-caa-glc and AB-caa-succ). [Due to multiple auxotrophies it is not possible to culture ppGpp^0^ cells in minimal medium without amino acid supplementation (Xiao et al., 1991)]. The ppGpp^0^ strain PAL3540 encodes loss-of-function mutations in both *relA* and *spoT* (MG1655 *spoT∷cat, relA∷kan)*, and is thus unable to synthesize the alarmone.

Mean square area remained inversely proportional to growth rate in ppGpp^0^ cells, indicating that size and nutrient availability remain coupled in the absence of ppGpp (Figure 4B; Table S2). Larger than congenic wild-type cells—potentially due to “free running” fatty acid synthesis in the absence of the alarmone—ppGpp^0^ mutants cultured in LB-glc and LB had mean square areas of *a* = 6.25 μm^2^ (*l* = 4.46 μm, *w* = 1.16 μm, τ τ 1.92 d/h) and 7.23 μm^2^ (*l* = 6.19 μm, *w* = 1.14 μm, τ = 1.68 d/h) respectively, while those cultured in AB-caa-glc had a mean square area of 3.01 μm^2^ (*l* = 2.98 μm, *w* = 0.97 μm, τ = 1.12 d/h) and those in AB-caa-succ a mean square area of 3.46 μm^2^ (*l* = 3.45 μm, *w* = 1.00 μm, τ = 0.96 d/h) (Figure 4B; Table S2). While we observed some extremely long ppGpp^0^ cells during growth in LB and LB-glc, such cells were absent during growth in minimal medium (Figure S3).

### ppGpp is required to maintain cell envelope integrity in the absence of fatty acid synthesis

Defects in fatty acid synthesis are incompatible with reductions in ppGpp synthesis (Yao et al., 2012a). Although not essential in a wild-type background, *fabH* is absolutely required in strains that are defective in one or both ppGpp synthases, RelA and SpoT (Yao et al., 2012a). Given this relationship, we wondered if an increase in intracellular [ppGpp] in response to defects in fatty acid synthesis might be protective, down-regulating other aspects of biosynthetic capacity to ensure that accumulation of cytoplasmic material does not exceed the capacity of the plasma membrane. To test this possibility, we examined the viability and membrane integrity of wild-type and ppGpp^0^ cells cultured in the presence of 500 μg/ml cerulenin (~5 times the MIC for wild-type cells).

Supporting a protective role for the alarmone, ppGpp^0^ cells were four orders of magnitude more sensitive to treatment with the fatty acid inhibitor cerulenin than their wild-type counterparts (Figure 5A). Cerulenin is generally viewed as bacteriostatic antibiotic, blocking synthesis of fatty acids without significantly impairing viability. Consistent with bacteriostatic activity, the plating efficiency of wild-type *E. coli* (MG1655) cells was largely unchanged after 4 hours of growth in the presence of 500 μg/ml cerulenin. In contrast, cerulenin treatment of ppGpp^0^ cells was bactericidal with plating efficiency dropping 10,000-fold over the course of 2.5 hours (Figure 5A).

**Figure 5.**
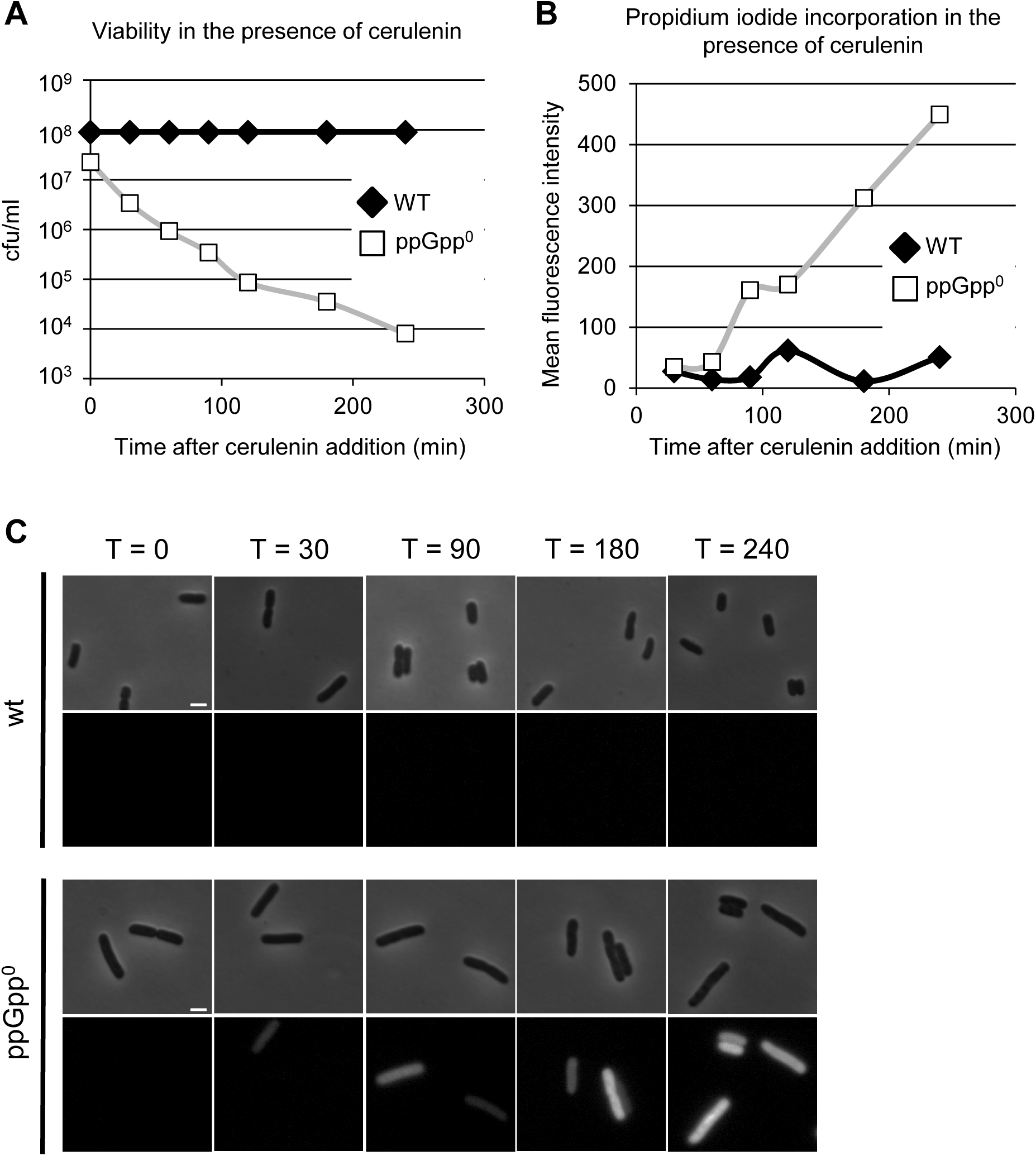
ppGpp is required to preserve membrane integrity and cell viability in the absence of fatty acid synthesis. (A) Colony forming units versus time for wild-type MG1655 and ppGpp^0^ strains. Cells were grown to OD_600_ of ~0.4 in LB-glc, treated with 500 μg/ml of cerulenin (at t = 0) and plated on LB agar every 30 min following cerulenin addition for 240 min. (B) Mean fluorescence intensity versus time of wild-type and ppGpp^0^ cells treated with cerulenin in the presence of the nucleic acid stain propridium iodide (diameter >2nm) which is unable to efficiently cross intact cellular membranes. (C) Images of cells sampled at selected time points from (B). Intense fluorescence is indicative of loss of membrane integrity. Scale bar = 2μm.

To assess the impact of cerulenin on cell envelope integrity, we visualized uptake of the dye, propidium iodide (PI), in wild-type and ppGpp^0^ cells in real time following cerulenin addition. PI is a nucleic acid intercalating agent that is unable to efficiently cross intact cellular membranes. When bound to nucleic acids, the fluorescence intensity of PI increases 20 to 30-fold. Due to its large size, however, PI requires pores of at least 2 nm to enter cells and bind to nucleic acid, a size generally incompatible with viability. PI fluorescence is thus a good indicator of loss of membrane integrity (Arndt-Jovin and Jovin, 1989; Bowman et al., 2010; Golzio et al., 2002).

PI staining strongly supports the idea that ppGpp accumulation preserves membrane integrity in response to inhibition of fatty acid synthesis through down regulation of other aspects of macromolecular synthesis (Figure 5B and C). The entirety of the ppGpp^0^ population exhibited bright red fluorescence indicative of a severe loss of membrane integrity three hours after the addition of 500 μg/ml of cerulenin (Figure 5C). In contrast, only a few PI-positive cells were visible in wild-type cells at the same time point. Calculation of mean fluorescence intensity suggests that wild-type cells were largely impermeable to the dye: 28 AU/cell (AU= arbitrary units) 30 minutes after the addition of drug rising to only 51 AU/cell two hours after the addition of drug. In contrast, ppGpp^0^ cells readily incorporated PI after the addition of cerulenin The mean fluorescence intensity of ppGpp^0^ mutants reached ~50 AU/cell within 60 minutes of cerulenin treatment and rose to 450AU/cell at 2.5 hours (Figure 5B).

## Discussion

This study firmly establishes lipid availability, and by extension cell membrane capacity, as a primary biosynthetic determinant of *E. coli* cell size (Figure 6). Increasing lipid synthesis, via overproduction of the transcriptional activator FadR, led to dose-dependent increases in cell size, while inhibiting lipid synthesis led to dose-dependent reductions in cell size (Figures 1 and 3). The modest reduction in length resulting from treatment with the translation inhibitor chloramphenicol was abolished by the addition of oleic acid to the medium (Figure 2B), further reinforcing a direct link between cell size and lipid availability. Inhibiting transcription had no detectable impact on cell morphology.

**Figure 6.**
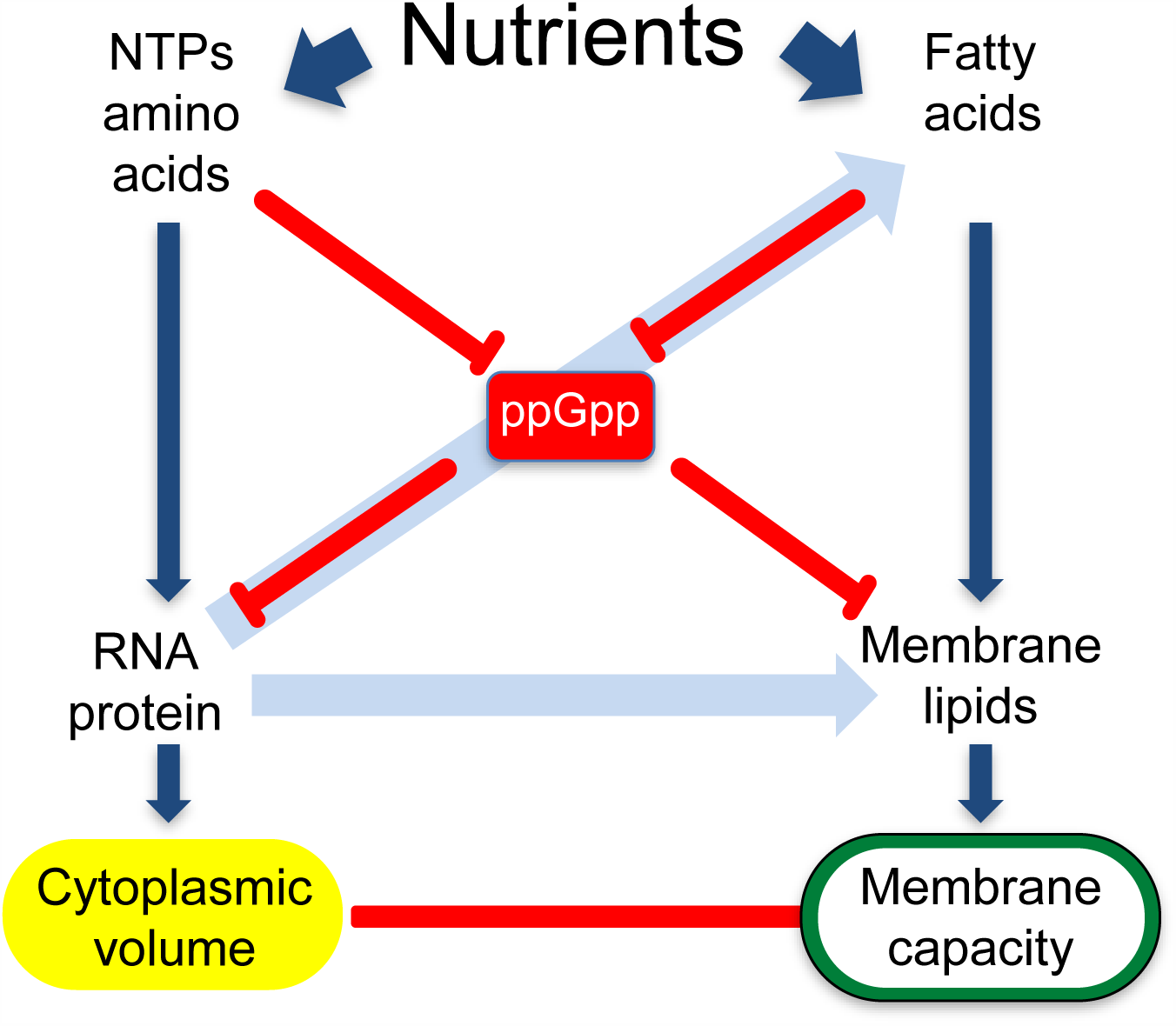
Plasma membrane capacity is coupled to cell volume via the alarmone ppGpp. During growth in nutrient rich medium, carbon-dependent flux (dark blue arrows) through fatty acid synthesis is high, increasing the pool of lipids available for cell envelope synthesis and with it cell size. In nutrient poor medium, flux through lipid synthesis is curtailed, reducing membrane capacity and cell volume. Synthesis of cytoplasmic material (e.g. RNA, protein) is similarly dependent on nutrient availability. RNA and protein synthesis impact synthesis of fatty acids and lipids indirectly (light blue arrows) via production of key enzymes. Lipid synthesis is coupled to synthesis of cytoplasmic material via the alarmone ppGpp (red lines).

A direct relationship between lipid synthesis and cell size provides a potential explanation for both the observation that bacterial cells add a constant volume (Δ) each generation under steady-state conditions as well as the strikingly linear relationship between nutrient availability and cell size. During balanced growth, a constant rate of flux through fatty acid synthesis coupled with a relatively invariant cell cycle clock translates into the addition of a constant volume of material in each generation. Shifting cells to carbon rich medium increases flux through fatty acid synthesis, transiently altering the balance between biosynthetic capacity and cell cycle progression until a new equilibrium is established albeit at a larger size (Δ). This model is supported both by data indicating that the rate of fatty acid synthesis is proportional to carbon availability (Davis et al., 2000; Li and Cronan, 1993) as well as Stephen Cooper’s observation that *E. coli* immediately increases its growth rate upon a shift to nutrient rich medium but delays division until cells achieve a new, larger size (Cooper, 1969). In *E. coli*, lipids are effectively all directed to the membrane (Parsons and Rock, 2013). Changes in the pool of lipids available for cell envelope biogenesis are thus expected to proportionally impact cell size.

Importantly, this study is the first to definitively place lipid synthesis downstream of ppGpp with regard to cell size. Inducing expression of the transcriptional activator of fatty acid synthesis, FadR, not only increased the size of wild-type cells, but also overcame the size defect associated with overproduction of the ppGpp synthase RelA (Figures 3 and 4A). Moreover, treatment with chloramphenicol, which mimics a “relaxed” (low ppGpp) state, reduced cell size in a lipid-dependent manner (Figure 2B). Cells defective in ppGpp production retained a linear relationship between size and nutrient availability, further reinforcing the idea that flux through lipid synthesis is a primary dictator of cell size independent of the alarmone (Figure 4B).

### Changes in lipid synthesis mimic the impact of nutrients on cell size and morphology

Like nutrients, altering flux through fatty acid synthesis impacted both the length and width of *E. coli* cells (Figures 1 and 3, Tables S2, S3 and S5). In our hands *E. coli* cultured in nutrient poor medium were ~30% shorter and ~25% thinner than their counterparts cultured in nutrient rich medium (Figure 1; Table S2). Cells cultured in nutrient rich LB-glc with 70 μg/ml of cerulenin exhibited a 20% reduction in length and a 10% reduction in width, in total a 30% reduction in square area that rendered them essentially indistinguishable from their counterparts cultured in AB-caa-glc. Overproduction of FadR increased cell length by 60% and cell width by 15%, increasing square area ~2-fold, a morphological change proportional to that observed in wild-type *E. coli* cells shifted from AB-succ to LB-glc [Figure 1 (Carbon); Figure 4; Tables S2 and S5].

The significant impact of lipid synthesis on both cell length and cell width distinguishes it from the relationship between cell cycle progression and cell size. Inhibiting divisome assembly can dramatically increase cell length, however, cell width is largely unaffected and in some cases may even be reduced (Harris and Theriot, 2016; Young, 2010). Delaying DNA replication initiation by depleting DnaA similarly leads to significant increases in cell length but does not detectably impact width (Pritchard et al., 1978).

### ppGpp: a master regulator coordinating lipid synthesis with other aspects of anabolic metabolism to preserve cellular integrity

A role for plasma membrane capacity as a primary determinant of cell size has far reaching implications for essentially all major aspects of cellular physiology that are sensitive to nutrient availability. Most obviously, lipid synthesis must be coordinated with synthesis of cytoplasmic material—particularly DNA, RNA and protein—to link surface area expansion with cytoplasmic volume and maintain cell envelope integrity. In this regard, our data suggest ppGpp plays a critical role, ensuring that flux through fatty acid synthesis is coupled to other aspects of biosynthetic capacity (Figures 5 and 6). In contrast to wild-type cells for which the addition of the fatty acid synthesis inhibitor cerulenin was bacteriostatic, ppGpp^0^ mutants exhibited a 10-fold reduction in viability within 30 min of addition of cerulenin, and gradually became permeable to propidium iodide (Figure 5). A role for ppGpp in coordinating fatty acid synthesis with cytoplasmic aspects of biosynthetic capacity is consistent with the longstanding observation that defects in ppGpp synthesis impair the ability of cells to respond to nutrient downshift (Murray et al., 2003; Ross et al., 2013; Sarubbi et al., 1988).

Loss of cell envelope integrity may also be a consequence of defects in coordination between lipid production and cell wall synthesis. Accumulation of ppGpp leads to down regulation of enzymes required for peptidoglycan synthesis and reduces the rate of diaminopimelic acid incorporation (Ishiguro and Ramey, 1976, 1978; Traxler et al., 2008). A negative relationship between ppGpp accumulation and cell wall synthesis is consistent with both our findings and the recent suggestion from Harris and Theriot that surface growth is coordinated with cell volume via growth rate-dependent changes in accumulation of cell wall precursors (Harris and Theriot, 2016). Independent of ppGpp, the reliance of peptidoglycan synthesis on availability of the 55-carbon lipid carrier, undecaprenyl pyrophosphate, may render it intrinsically dependent on flux through central carbon metabolism. In *B. subtilis* lipid production is enhanced upon depletion of enzymes required for synthesis of peptidoglycan precursors, suggesting the potential for a feedback loop coupling peptidoglycan and lipid synthesis to one another (Mercier et al., 2013).

Although unlikely to directly impact cell envelope integrity, membrane synthesis must also be synchronized with cell cycle progression to ensure the production of viable daughter cells. ppGpp has previously been implicated in coordinating amino acid starvation with both DNA replication and chromosome segregation. In *E. coli*, treatment with serine hydroxamate results in ppGpp-dependent inhibition of replication elongation in addition to a reduction in cell size (Tehranchi et al., 2010). ppGpp is also reported to interfere with chromosome segregation, although the mechanism underlying this phenomenon in unclear (Ferullo and Lovett, 2008). Induction of ppGpp upon inhibition of fatty acid synthesis serves an adaptive function for the alphaproteobacterium *C. crescentus*, down-regulating production of both the global cell cycle regulator CtrA as well as DnaA to prevent DNA replication in the absence of growth (Leslie et al., 2015; Stott et al., 2015).

### Beyond *E. coli*

It remains to be seen if lipid synthesis and plasma membrane capacity are widely conserved as a link between nutrient availability and cell size. Little is known about the relationship between fatty acid synthesis and ppGpp synthesis in the Gram-positive model bacterium *B. subtilis.* Moreover, many bacteria, including the pathogen *Mycobacterium tuberculosis*, store lipids in self-contained, cytoplasmic vacuoles. How lipid storage is balanced with cell envelope biogenesis in such organisms remains an open question. Further afield, the target of rapamycin (TOR) kinase, promotes the growth and proliferation of eukaryotic cells in response to increases in nutrient availability, including the presence of growth factors and other generally auspicious environmental conditions (Laplante and Sabatini, 2009; Loewith and Hall, 2011). While much of the literature has focused on the role of TOR in regulating protein synthesis, TOR has also been implicated in stimulating lipid synthesis, both for cell membrane and storage purposes (Laplante and Sabatini, 2009). It is thus of great interest to know whether the relationship between lipid pools and cell size that we observed in *E. coli*, holds true in eukaryotes, where fatty acid synthesis is restricted to the mitochondria.

## Materials and Methods

### Bacterial strains and culture conditions

*E. coli* strains and plasmids are listed in table S1. Standard techniques were employed for P1 vir transductions and other genetic manipulations. Cells were cultured in lysogeny broth (LB) + 0.2% glucose or AB minimal medium (Clark, 1967) supplemented with 10 μg/ml thiamine, 5 μg/ml thymidine and 0.5% casamino acids, 0.2% glucose or 0.4% succinate as indicated. Strains carrying plasmids were cultured with 100 μg/ml ampicillin or 50 μg/ml kanamycin as required. Cells were used for experimentation at early exponential growth phase (OD_600_ = 0.15-0.3) unless stated otherwise. To achieve balanced growth, cells were cultured at 37°C in the indicated medium from a single colony to early exponential phase with vigorous shaking (200 rpm). Cultures were then back-diluted into fresh medium to an OD_600_ of 0.005 and cultured to early exponential phase prior to being sampled and fixed for analysis. For experiments in minimal medium, cells were first cultured from a single colony overnight prior to being back-diluted into minimal medium to an OD_600_ of 0.005 and subjected to an additional round of growth and back-dilution as above. In experiments utilizing sub-inhibitory concentrations of rifampicin, chloramphenicol or cerulenin to inhibit RNA, protein or fatty acid synthesis, antibiotics were added when cultures were back-diluted. In experiments involving isopropyl β-D-1- thiogalactopyranoside (IPTG) or exogenous fatty acids, they were added along with antibiotics at the indicated concentrations. Mass doubling times were calculated from OD600 readings taken every 30 minutes using the online doubling time calculator, cell calculator++ (Roth, 2006). For experiments utilizing the ppGpp^0^ strain, PAL3540 (MG1655 *relA∷kan spoT∷cat)*, cells were plated on AB minimal medium + 0.2% glucose and LB agar to confirm that they had not accumulated suppressor mutations that suppressed the auxotrophies associated with the absence of ppGpp.

### Microscopy and image analysis

Images were acquired from samples on 1% agarose (in PBS) pads with an Olympus BX51 microscope equipped with a 100X Plan N (N.A. = 1.25) Ph3 objective (Olympus), X-Cite 120 LED light source (Lumen Dynamics), and an OrcaERG CCDcamera (Hammamatsu Photonics, Bridgewater, N.J.). Filter sets for fluorescence were purchased from Chroma Technology Corporation. Nikon Elements software (Nikon Instruments, Inc.) was used for image capture. Cells were fixed with 2.6% paraformaldehyde + 0.008% glutaraldehyde, and stained with the membrane stain FM 4-64_fx_ (ThermoScientific) prior to imaging as described previously (Levin, 2002). Cell size was determined from phase contrast images with the ImageJ pulgin Coli-Inspector (Vischer). FM 4-64 membrane stain was employed to differentiate between cells that had completed division and those that had not. For analysis of cell cycle progression, cells were considered positive for FtsZ-ring formation if they contained a visible band or two spots of GFP-FtsZ fluorescence at midcell. Cells were also considered positive for FtsZ-ring formation if a single spot of GFP-FtsZ was visible at the midpoint of an invaginating septum.

### Measurement of steady-state intracellular nucleotides by thin layer chromatography

Strains were grown from single colonies in low-phosphate MOPS (Teknova) at 37°C with shaking (1X MOPS, 1 mM KH2PO4, 0.2% glucose, 0.4% casamino acids) and appropriate antibiotics to early logarithmic phase, then diluted to OD_600_ = 0.005, at which time ^32^P-orthophosphate was introduced to label the cells. Once cells reached early logarithmic phase again, samples were collected for nucleotide extraction using 2M ice cold formic acid. Nucleotides were measured using thin layer chromatography and quantified using ImageQuant software (Molecular Dynamics) as described previously (Wang et al., 2007). Intensities were normalized to the number of phosphates in the corresponding nucleotide. ppGpp levels are normalized to GTP level at steady state with signals from steady-state (p)ppGpp^0^ cells subtracted.

### Cell viability assays

Cells were grown in LB-0.2% glucose at 37°C from a single colony to OD_600_ = 0.2-0.3, back-diluted to OD_600_ = 0.005, and returned to 37°C with shaking. At OD_600_ ~ 0.4, cultures were divided equally and exposed to either 500 μg/ml cerulenin, an equal volume of 95% EtOH as a solvent control or left untreated. Serial dilutions of each sample were spread on LB agar every 30 minutes after treatment. Colony forming units (CFUs) were enumerated after 24 h of incubation at 37°C. Because ppGpp^0^ cells tend to accumulate suppressor mutations in RNA polymerase, ppGpp^0^ samples were plated on AB-minimal + 0.2% glucose and LB agar to identify suppressor mutants. The frequency of ppGpp^0^ suppressor mutants was calculated as the ratio of CFU/ml recovered on minimal vs. LB plates. Data was only included from experiments where the ratio of CFU/ml recovered on minimal vs. LB plates was less than 1:10.

### Assessment of cell envelope integrity via propidium iodide incorporation

Cells were grown in LB-0.2% glucose at 37°C from a single colony to OD_600_ = 0.2-0.3, back-diluted to OD_600_ = 0.005, and returned to 37°C. At OD_600_ ~ 0.4, cultures were divided equally and exposed to either 1.5 μM propidium iodide alone, or propidium iodide and either 500 μg/ml cerulenin or an equal volume of 95% EtOH as a solvent control. At indicated intervals, samples were removed and mounted on 1% agarose pads for phase contrast and fluorescence imaging. The extent of propidium iodide incorporation was measured by quantitative fluorescence microscopy. Briefly, fluorescence images were background corrected, bacteria from phase contrast images were outlined as regions of interest (ROIs) and the average fluorescence intensity per cell of the ROIs was determined from background corrected fluorescence images.

### Statistical analysis

A minimum of three independent biological replicates were performed for each experimental data set. Data were expressed as means ± standard errors of the means (SEM). *P* values were calculated using a standard two-tailed Student’s *t* test. Asterisks indicate a significant difference between the results for indicated experimental conditions (*, *P* < 0.05; **, *P* < 0.01).

## Author Contributions

S.V., P.A.L., J.D.W., and J.L.T. designed the experiments. S.V. and J.L.T. performed the experiments. P.A.L. and S.V. wrote the manuscript.

## Acknowledgements

We are indebted to Mike Cashel, John Cronan, Christophe Herman, Rick Gourse, Charles Rock, and Fuzhong Zhang for the kind gift of *E. coli* strains and plasmids, and to Dianne Duncan, Norbert Vischer and Anne Katherine Fee for assistance with image capture and analysis. We would also like to thank Gayle Bentley, Natacha Ruiz and members of the Levin and Zaher labs for numerous discussions about matters technical and philosophical over the course of this research, and Heidi Arjes, Rick Gourse, Daniel Haeusser and Suckjoon Jun for critical reading of the manuscript. This work was supported by National Institutes of Health grants GM64671 to PAL and GM084003 to JDW.

